# Recent outbreaks of chikungunya virus (CHIKV) in Africa and Asia are driven by a variant carrying mutations associated with increased fitness for *Aedes aegypti*

**DOI:** 10.1101/373316

**Authors:** Irina Maljkovic Berry, Fredrick Eyase, Simon Pollett, Samson Limbaso Konongoi, Katherine Figueroa, Victor Ofula, Helen Koka, Edith Koskei, Albert Nyunja, James D. Mancuso, Richard G. Jarman, Rosemary Sang

## Abstract

**Background:** In 2016, a chikungunya virus (CHIKV) outbreak was reported in Mandera, Kenya. This was the first major CHIKV outbreak in the country since the global re-emergence of this virus, which arose as an initial outbreak in Kenya in 2004. Therefore, we collected samples and sequenced viral genomes from the 2016 Mandera outbreak.

**Methodology/Principal Findings:** All Kenyan genomes contained two mutations, E1:K211E and E2:V264A, recently reported to have an association with increased infectivity, dissemination and transmission in the *Aedes aegypti* (*Ae. aegypti*) vector. Phylogeographic inference of temporal and spatial virus relationships using Bayesian approaches showed that this *Ae. aegypti* adapted strain emerged within the East, Central, and South African (ECSA) lineage of CHIKV between 2005 and 2008, most probably in India. It was also in India where the first large outbreak caused by this strain appeared, in New Delhi, 2010. More importantly, our results also showed that this strain is no longer contained to India, and that it has more recently caused several major outbreaks of CHIKV, including the 2016 outbreaks in India, Pakistan and Kenya, and the 2017 outbreak in Bangladesh. In addition to its capability to cause large outbreaks in different regions of the world, this CHIKV strain has the capacity to replace less adapted wild type strains in *Ae. aegypti-*rich regions. Indeed, all the latest full CHIKV genomes of the ECSA Indian Ocean Lineage (IOL), from the regions of high *Ae. aegypti* prevalence, carry these two mutations, including samples collected in Japan, Australia, and China.

**Conclusions/Significance:** Our results point to the importance of continued genomic-based surveillance of this strain’s global spread, and they prompt urgent vector competence studies in Asian and African countries, in order to assess the level of vector receptiveness, virus transmission, and the impact this might have on this strain’s ability to cause major outbreaks.

**Author summary:** Chikungunya virus (CHIKV) causes a debilitating infection with high fever, intense muscle and bone pain, rash, nausea, vomiting and headaches, and persistent and/or recurrent joint pains for months or years after contracting the virus. CHIKV is spread by two mosquito vectors, *Aedes albopictus* and *Aedes aegypti*, with increased presence around the globe. In this study, we report global spread of a CHIKV strain that carries two mutations that have been suggested to increase this virus’ ability to infect the *Aedes aegypti* mosquito, as well as to increase CHIKV’s ability to be transmitted by this vector. We show that this strain appeared sometime between 2005 and 2008, most probably in India, and has now spread to Africa, Asia, and Australia. We show that this strain is capable of driving large outbreaks of CHIKV in the human population, causing recent major outbreaks in Kenya, Pakistan, India and Bangladesh. Thus, our results stress the importance of monitoring this strain’s global spread, as well as the need of improved vector control strategies in the areas of *Aedes aegypti* prevalence.

## Introduction

In May 2016, the Kenyan Ministry of Health (KMoH) reported an outbreak of chikungunya virus (CHIKV) in Mandera County on the border with Somalia. During this time in Somalia, outbreaks of CHIKV were occurring in the neighboring Bula Hawa region, originating from Mogadishu. In Mandera town, 1,792 cases were detected, and an estimated 50% of the health work force was affected by this virus. A cross-border joint response was coordinated between Kenya and Somalia to control the outbreak (*1*). This was the first reported outbreak of CHIKV in Kenya since 2004.

The previous large CHIKV outbreak in Kenya occurred in Lamu Island in 2004, with an estimated 75% of the population infected (*2*). The disease also spread to the coastal city of Mombasa by the end of 2004, and further to the Comoros and La Réunion islands, causing large outbreaks in 2005-2006. On La Réunion island, unusual clinical complications were reported in association with CHIKV infection, and viral isolate sequences revealed presence of an alanine-to-valine mutation in the E1 glycoprotein at position 226 (E1:A226V) (*3*). This specific mutation was shown to confer enhanced infection of the *Aedes albopictus* vector, which was also the dominant mosquito species suspected to be responsible for the transmission of CHIKV on the island of La Réunion (*4, 5*).

Since then, the E1:A226V mutation has been observed in many of the genomes in the CHIKV lineage spreading in the East, Central, and South African region (ECSA lineage), and has been shown to have emerged through convergent evolution in at least four different occasions (*6*). This remarkable emergence and spread of CHIKV adaptation to the *Ae. albopictus* vector prompted additional studies on the genetic plasticity of this virus, looking for additional biomarkers associated with virus transmission capacity, fitness and pathogenicity. In 2009, a mutation E2:L210Q was observed in CHIKV isolates from Kerala, India, and was shown to further increase the E1:A226V enhancement of the CHIKV infection in *Ae. albopictus* (*7*). Other mutations have also been described with the ability to enhance infection in this vector, information of increased importance as *Ae. albopictus* is rapidly expanding throughout the world (*8*). The *Ae. albopictus* mosquito is believed to have originated in Asia, and is today most commonly found on the east coasts of the Americas and east Asia, with some prevalence in Southeast Asia, India and Africa, and with increased spread to regions with lower temperatures, such as southern Europe, southern Brazil, northern China and northern United States (*8*).

Another important vector of CHIKV is *Aedes aegypti,* a container-breeding, domesticated mosquito mainly found in urban areas and feeding largely on humans during daytime hours. *Ae. aegypti* originated from the ancestral zoophilic *Ae*. *aegypti formosus* native to Africa. *Ae. aegypti* now is most common in tropical and sub-tropical regions, such as Africa, India, Southeast Asia and Australia (*8*). Recently, two mutations, E1:K211E and E2:V264A, have been reported in the CHIKV to be associated with increased adaptation to *Ae. aegypti* vectors (*9*). These two mutations, in the background of the wildtype E1:226A, are believed to increase virus infectivity, dissemination and transmission in *Ae. aegypti,* with no impact on virus fitness for the *Ae. albopictus* vector (*6, 9*). E1:K211E was first observed in genomes sampled in 2006 from the Kerala and Puducherry regions, India, and simultaneous presence of both mutations was first observed in 2009-2010 in Tamil Nadu and Andhra Pradesh, India (*10, 11*).

In Kenya, the predominant CHIKV vector is *Ae. aegypti*, and local vector competence studies have shown it is capable of transmitting the virus in this region (*12*). Given the recent large outbreak of CHIKV in the rural setting of Mandera County, Kenya, we analyzed CHIKV genomes sequenced from this outbreak for presence of adaptive mutations associated with both *Ae. albopictus* and *Ae. aegypti*. Along with estimating time and origins of the Mandera CHIKV outbreak, we investigated the origins and the time of emergence of its detected vector adaptive mutations. Our results point to an increased spread of the *Ae. aegypti* adapted CHIKV strain capable of causing new large outbreaks in different regions of the world.

## Methods

### Ethics statement

The study was carried out on a protocol approved by the Walter Reed Army Institute of Research (WRAIR #2189) and Kenya Medical Research Institute’s Scientific and Ethics Review Unit (SERU#3035) as an overarching protocol guiding investigation and reporting of arbovirus/Hemorrhagic fever outbreaks in Kenya. Since the blood samples were collected from an outbreak and no human data was collected, this was deemed to be a non-human research study and no consent was required. The protocol was approved for additional analysis of outbreak samples and for publication of results. STROBE checklist is in the supplementary file S1_Checklist.

### Samples and sequencing

From May 2016, following reports of widespread incidence of febrile illness with severe joint pains in Mandera (a city at the border with Somalia) and its environs, samples were collected from suspected cases of all ages and sex, using standard practices by the Kenya Ministry of Health staff. The case definition used was “any patient presenting with sudden onset of fever >38.5 °C, with severe joint/muscle pains and headache within the last 3-5 days within Mandera County, the person should either be a resident of or visiting Mandera”. Chikungunya infection was confirmed by CHIKV specific RT-PCR and partial genome Sanger sequencing at the Kenya Medical Research Institute (KEMRI) laboratories. Vero cell culture inoculations were performed on the samples to obtain isolates for in-depth studies. For method details on infection confirmation, virus isolation and Sanger sequencing see S2_Appendix. A subset of chikungunya-confirmed positive samples from acutely ill patients were further subjected to high throughput full genome sequencing at the KEMRI laboratories. The prepared cDNA was quantified using Qubit 3.0 Fluorimeter and the dsDNA HS Assay Kit (Life Technologies). The complimentary DNA was fragmented enzymatically and tagged using the Nextera XT DNA Library kit (Illumina, San Diego, CA, USA). Each sample was assigned a unique barcode sequence using the Sequence libraries prepared with the Nextera-XT kit (Illumina) and sequenced on the MiSeq platform according to manufacturer’s instructions (Illumina). Ten full genomes and five partial genomes were assembled by combining both de novo and reference mapping assemblies. De novo assemblies were performed using Abyss (*13*) and Trinity (*14*), and reference mapping was performed using ngs_mapper (*15*). Sequences have been submitted to GenBank under accession numbers MH423797-MH423811.

### Phylogenetic analyses and selection

A full genome CHKV reference dataset was downloaded from GenBank and curated in TempEst (*16*). All outlier genomes were removed and the cleaned genomes (n=466) were aligned to the ten assembled full genomes from Mandera, Kenya, using MEGAv7 (*17*). This large dataset was scanned for presence of recombination using Recombination Detection Program version 4 (RDP4) (*18*), with a minimum of 3 methods with significant (p<0.05) recombination signal required to call a genome recombinant. Maximum Likelihood trees were inferred using PhyML (*19*) and RaxML (*20*), with GTR+G+I model of evolution, as determined by jModelTest2 (*21*). Node confidence values were derived by aLRT (PhyML) and bootstrap of 500 (RaxML). After evaluating the temporal structure using TempEst, the trees were used for an informed down-sampling of the dataset. The down-sampled dataset (ECSA lineage only, n=115) was analyzed using the BEAST package (*22*) with GTR+G+I model of substitution, bayesian skyline plot, relaxed lognormal molecular clock, asymmetrical trait (location) distribution. BEAST was run for 800 million generations, with subsampling every 80,000 and 10% burn-in. The Maximum Clade Credibility (MCC) tree was summarized using TreeAnnotator. Amino acid mutations at each node for the resulting trees were mapped using TreeTime (*23*). Because viral genomes from several additional recent CHIKV outbreaks form India and Bangladesh were only sequenced in their E1 gene, we also inferred ML tree of the partial E1 region including these sequences, using PhyML and TN93+G+I model of evolution, as determined by jModelTest2. Selection analyses were performed using the HyPhy package (*24*). Presence of site specific selection was determined by a likelihood approach using FEL and by Bayesian method FUBAR with probability level threshold of 0.95 (*25, 26*). Episodic selection was investigated using MEME (*27*), and any selection acting on tree branches was determined by aBSREL (*28*). Selection analyses were performed on the large full genome dataset used for the ML trees (ECSA only) and the small dataset used for BEAST analyses. FUBAR was performed on separated structural and non-structural genes from both datasets because of their large sizes.

## Results

### CHIKV causing the 2016 outbreak in Mandera, Kenya, was introduced in 2015

A total of 15 samples from the CHIKV outbreak in Mandera, Kenya, collected in May and June 2016, were sequenced. Ten of the sequenced samples produced full CHIKV genomes, and five of the samples had partial genomes. No recombination was detected in the genomes. ML trees inferred by PhyML and RaxML were concordant and showed that all Kenyan genomes belonged to the ECSA lineage of CHIKV. Consistently with their outbreak origins, the genomes from Mandera clustered in a monophyletic clade defined by very short branches, indicating limited genetic diversity (Figure 1). The Kenyan cluster was most closely related to genomes from Japan and India; however, the long branch leading to this cluster indicated a probable lack of sampling of viruses more related to the Kenyan outbreak. Two genomes from the CHIKV outbreak in Kenya in 2004 were located more basally in the tree and were very different from the virus causing the outbreak of 2016. This indicated that the 2016 outbreak virus did not directly evolve from a CHIKV circulating in Kenya since 2004, but was introduced at a later time point. The most recent common ancestor (MRCA) of the Kenyan 2016 viruses existed in mid-2015 [2015.6; HPD 95%=2015.1-2015.9], making this the latest possible time point of introduction of this virus into Mandera, Kenya. The ancestor shared with the most closely related genomes from Japan and India existed in late-2009 [2009.9; 95% HPD=2008.6-2011.5] and was estimated to have originated in India. These results indicate that the CHIKV causing the 2016 outbreak in Kenya was introduced into this region sometime by 2015, originating from a virus that existed in India in 2009. However, India might not have been the direct source of the CHIKV introduction into Kenya and more sampling is needed to, with greater precision, determine the exact origin of this introduction.

**Figure 1.**
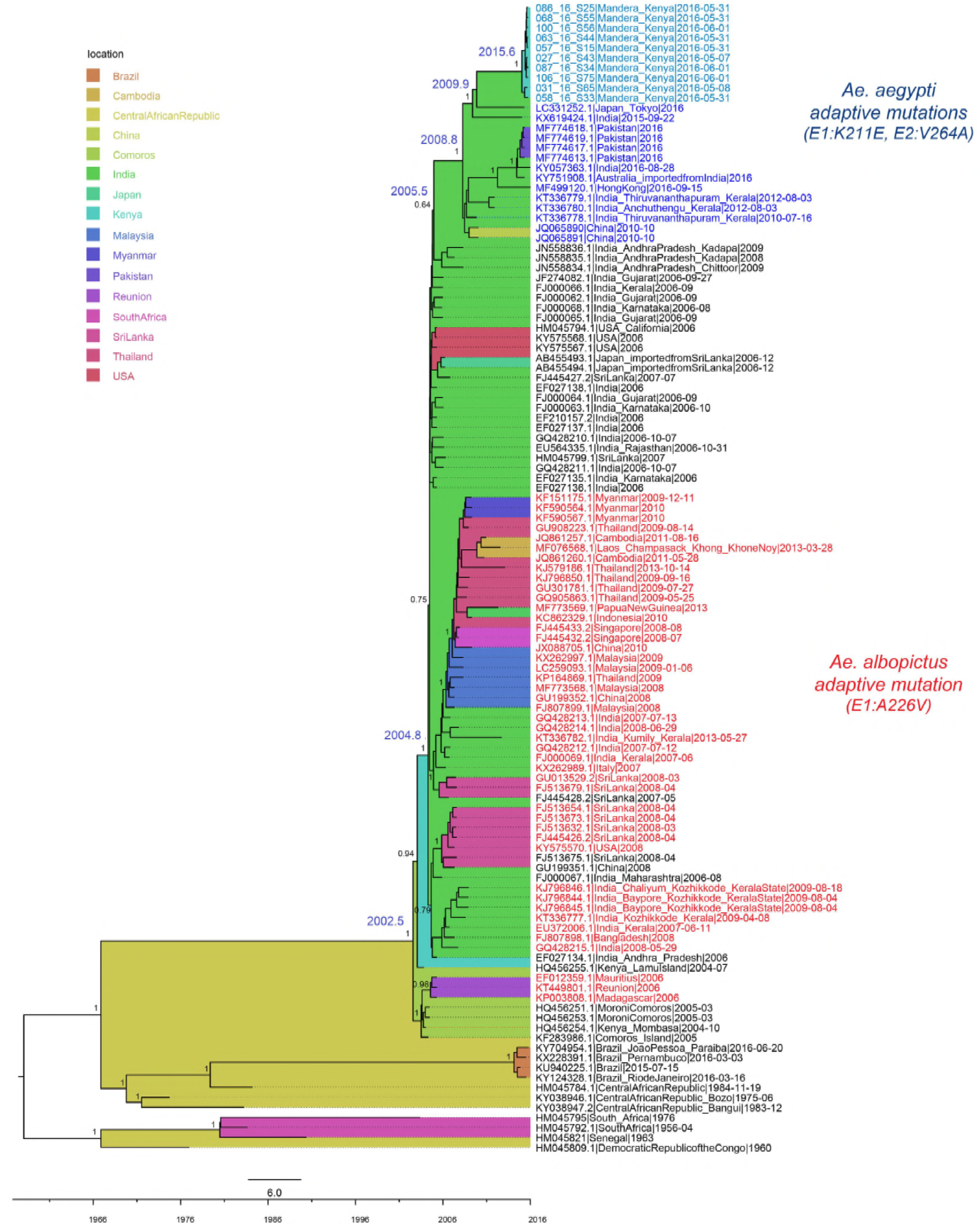
Full genome MCC tree of the CHIKV ECSA lineage. Estimated location origin is marked in colored tree background, according to the legend. Taxa in red text represent genomes containing the *Ae. albopictus* adaptive E1:A226V mutation, while all other taxa contain the wildtype. Taxa in blue (light blue for Kenya) represent genomes with the *Ae. aegypti* adaptive E1:K211E and E2:V264A mutations. Most important node supports are shown, as well as the estimated TMRCAs of nodes of interest.

### Mutations associated with increased fitness to Ae. aegypti emerged by 2008

Careful analyses of amino acid mutations previously suggested to associate with vector adaptation revealed that all Kenyan viruses contained two mutations (E1:K211E and E2:V264A) that, in the background of E1:226A, have recently been correlated with enhancement of CHIKV fitness for the *Ae. aegypti* vector (Table 1). All Kenyan genomes also had the E1:226A background amino acid. Further investigation of amino acids from the genomes surrounding the Kenyan samples in both the ML and MCC phylogenetic trees revealed a cluster of genomes, sampled from various regions of the world (Asia, Africa, Australia), also containing the two *Ae. aegypti* adaptive mutations in the background of E1:226A (Figure 1). These two amino acid changes were the only ones that characterized this cluster.

**Table 1.** Changes in positions previously associated with vector competence and pathology of CHIKV. Amino acids associated with each phenotype are shown in bold underlined font. Only available full genomes from each lineage were compared.

| Protein | Amino acid substitution | Phenotype | Asian lineage | ECSA lineage without *Ae.aegypti* adapted cluster | ECSA *Ae.aegypti* adapted cluster | Kenya |
| --- | --- | --- | --- | --- | --- | --- |
| E1 | A98T (6, 29) | Epistatic constraint on E1:A226V | <u>**T**</u> | A | A | A |
| E1 | A226V (4) | Enhanced infection of *Ae. albopictus* | A | A, <u>**V**</u> | A | A |
| E1 | K211E (9) | Enhanced fitness in *Ae. aegypti* in background of E1:226A | <u>**E**</u> | K, T, N | <u>**E**</u> | <u>**E**</u> |
| E2 | V264A (9) |  | V | V | <u>**A**</u> | <u>**A**</u> |
| E2 | R198Q (6) | Enhanced infection of *Ae. albopictus*, synergistic with E3:18F in background of E1:226V | R | R, <u>**Q**</u> | R | R |
| E2 | L210Q (6, 7) | Enhanced infection of *Ae. albopictus*, secondary to E2:A226V | L | L, <u>**Q**</u> | L | L |
| E2 | K233E/Q (6) | Enhanced infection of *Ae. albopictus* | K | K, <u>**E**</u> | K | K |
| E2 | K234E (6) | Enhanced infection of *Ae. albopictus* | K | K | K | K |
| E2 | K252Q (6) | Enhanced infection of *Ae. albopictus* – secondary mutation | K | K, <u>**Q**</u> | K | K |
| E3 | S18F (6) | Enhanced infection of *Ae. albopictus*, synergistic with E2:R198Q | S, <u>**F**</u> | S, <u>**F**</u> | S | S |
| nsP1 | G230R (30) | Increase replication in *Ae. albopictus* in combination with nsP3:524* | G, <u>**R**</u> | G, <u>**R**</u> | G | G |
|  | V326M (30) |  | V, <u>**M**</u> | V | V | V |
| nsP3 | *524R (31) | Attenuation of arthritis and pathology | *, L, C, <u>**R**</u> | *, C, <u>**R**</u> | * | * |

\*Opal STOP codon

The cluster containing viruses with the suggested *Ae. aegypti* adaptive mutations consisted of genomes from two different outbreaks, Kenya in 2016 and Pakistan in 2016, and additional genomes from India sampled between 2010 and 2016, and Japan, Hong Kong, Australia, all sampled in 2016. The MRCA of this cluster was estimated to have existed in India in late 2008 [2008.8; 95% HPD=2007.9-2009.5], indicating that the virus with dual E1:K211E and E2:V264A mutations emerged by the end of 2008 in this area of the world. The cluster shared a common ancestor with genomes from India, which did not contain the two *Ae. aegypti* adaptive mutations, in mid-2005 [2005.5; 95% HPD=2005.0-2006.1]. Although the node support for the ancestor of adapted and non-adapted strains was low, probably due to lack of sampling that would reveal the exact evolutionary relationships, it was estimated to have existed in India due to well supported ancestral nodes preceding this one. Thus, these results indicated that the *Ae. aegypti* adaptation most probably arose in India sometime between 2005 and 2008. The background E1:226A amino acid (non-red taxa, Figure 1) predominated in this part of the tree, while the *Ae. albopictus* adaptive E1:226V amino acid (red taxa, Figure 1) was mainly found in the sub-lineage containing genomes from Southeast Asia. Genomes from India sampled in years 2007-2009 were found containing E1:226A and E1:226V mutations, meaning that this country experienced simultaneous spread of both strains. The MRCA of all Indian viruses existed in 2004 [2004.8; 95% HPD=2004.3-2005.1], and seems to have originated from a virus related to the Kenyan Lamu Island outbreak. The 2004 genome from Mombasa, Kenya, was more related to the viruses from Comoros, Madagascar and La Réunion. The two Kenyan 2004 genomes shared a common ancestor that existed in 2002 [2002.5; 95% HPD=2001.6-2003.3] and was initially of a Central African Republic origin, however, the long branch indicates lack of sampling and makes it difficult to exactly determine viral geographical transmission during this time.

ML trees of the partial E1 region, including additional sequences from the CHIKV outbreaks in 2010 and 2016 from New Delhi, India, and from 2017 in Bangladesh, showed that genomes from these outbreaks also belonged to the *Ae. aegypti* adaptive cluster (Figure 2). Despite low phylogenetic signal due to the shorter E1 segment, the cluster was supported by high confidence value, 0.93. These results indicated that, in addition to the outbreaks of Pakistan and Kenya, the more recent outbreaks in India and Bangladesh were also most probably caused by the *Ae. aegypti* adapted strain.

**Figure 2.**
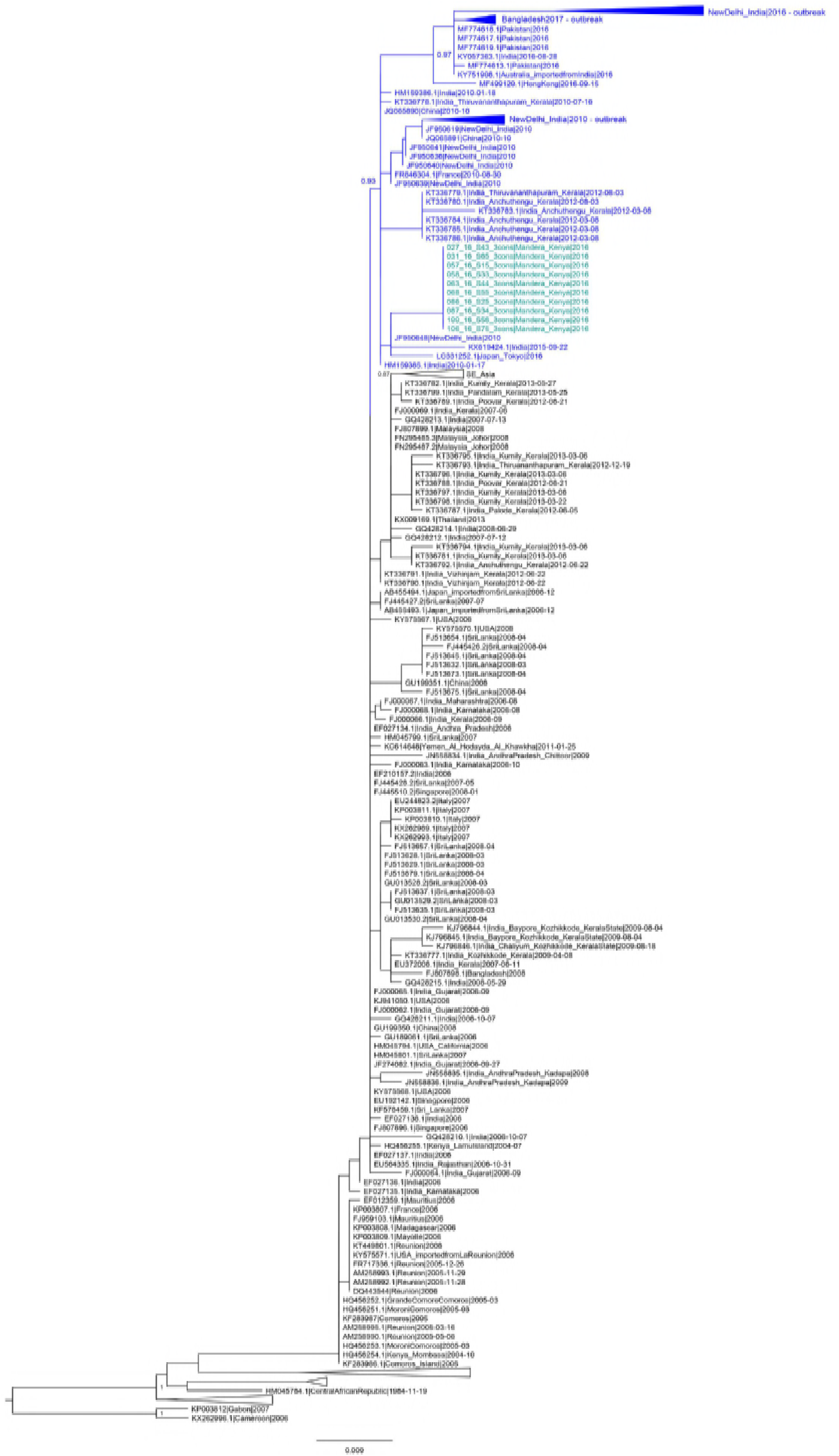
Partial E1 gene ML tree of the CHIKV ECSA lineage. Taxa in blue (light blue for Kenya) represent genomes with the *Ae. aegypti* adaptive E1:K211E mutation.

### Positive selection was acting on the E1:K211 but not the E2:V264A position

To investigate whether the E1:K211 and E2:V264A mutations appeared due to selective pressure and adaptation of the virus, we performed several tests for presence of positive selection, on both the large CHIKV dataset used for the ML trees, and on the smaller dataset used for BEAST analyses. Significant presence of positive selection in position E1:K211 was detected by both FUBAR (probability > 0.99), FEL (p≤0.05) and MEME (p<0.05) in all tested datasets. Position E2:V264A did not show any evidence of positive selection. No branch-specific selection was detected by aBSRL. Positions found to be under positive selection by all methods are listed in Table 2.

**Table 2.** Positively selected positions by method.

|  | FUBAR | FEL | MEME |
| --- | --- | --- | --- |
| ML dataset | E1:211 | nsP1:171, nsP3:117, E1:211 | nsP1:4, nsP1:82, nsP1:157, nsP1:171, nsP1:301, nsP1:407, nsP2:349, nsP2:457, nsP2:604, nsP3:117, nsP3:303, nsP4:81, nsP4:467, nsP4:605, E2:57, E2:178, 6K:47, E1:146, E1:211, E1:382 |
| **BEAST**<br>**dataset** | E1:211 | nsP1:171,<br>nsP3:117,<br>E1:211 | nsP1:4, nsP1:101, nsP1:171, nsP2:349, nsP3:117, nsP4:81,<br>nsP4:467, E2:57, 6K:47, E1:146, E1:211 |

## Discussion

Recent re-emergence and global spread of Chikungunya virus, coupled with its high morbidity and economic burden, has made it one of the more medically important arboviral diseases with major public health implications. CHIKV originated in Africa and the first known outbreak was recorded in today’s Tanzania in 1952-1953. Subsequently, sporadic outbreaks in Africa and larger epidemics in Asia were observed until the 1980s, followed by a period of decreased activity. The virus re-emerged in the early 2000s in Africa resulting in extensive and rapid spread throughout the world. In this study, we analyzed ten CHIKV genomes sampled from the 2016 outbreak in Mandera, Kenya, and compared them to the genomes from Kenya outbreak in 2004. We estimated the time of the 2016 outbreak emergence, as well as traced the emergence and movement of the *Ae. aegypti* adaptive mutations that characterized these genomes. Strains with increased vector fitness may have the potential of more efficient transmission resulting in large outbreaks, and may outcompete wild type viral variants in the regions with abundant vector populations.

Our results suggest that re-emerging CHIKV reached Kenya in mid-2002, where it split into two strains. One strain caused disease in Mombasa in 2004 and spread further to Comoros, Madagascar and La Réunion, where it obtained the *Ae. albopictus* adaptive mutation E1:226V. The other strain, still carrying the wild type E1:226A, spread north, causing the Lamu Island outbreak. Although the Lamu Island outbreak was observed first and was the largest in Kenya in 2004, current results suggest it did not seed the virus in Mombasa. Rather, the two shared a common ancestor that split into two different directions. Given that only one genome from each region is available, more data would be needed to completely resolve virus movement within Kenya during this period of time. Exact estimations of intercontinental routes of geographic spread are also limited by sampling skew, however, the wild type strain from Lamu Island was next observed causing outbreaks in 2005-2006 in India, mainly on the west and south of the country (Rajasthan, Gujarat, Maharashtra, Andhra Pradesh, Karnataka) (32-34). Shortly thereafter, in 2007, a strain carrying the *Ae. albopictus* adaptive E1:226V mutation was recorded in the province of Kerala (35). Both strains were found circulating in the following years in the country, between 2007 and 2013 (36, 37).

In India, both *Ae. albopictus* and *Ae. aegypti* are prevalent vectors, and the presence of the recently described *Ae. aegypti* adaptive mutation E1:K211E was first observed in the Kerala-Puducherry CHIKV outbreak of 2006 (*10*). Shortly thereafter, E1:K211E was accompanied by the second *Ae. aegypti* adaptive mutation, E2:V264A. This novel dual-mutation strain was first observed in Kerala in 2009, however, the Kerala-Puducherry CHIKV outbreak of 2006 was not sequenced in the E2 region (*11*). These results support our estimate that the *Ae. aegypti* adaptive strain probably arose in India sometime between 2005 and 2008. The first indication that this strain was capable of causing major outbreaks came from India in 2010, where it caused an outbreak in New Delhi (38). Following this, CHIKV continued to circulate in discrete regions throughout India, and in 2016 the adapted strain caused another large outbreak in New Delhi (39). Simultaneously, it also caused the outbreaks in Pakistan and, as reported here, in Kenya (40). The Kenyan outbreak was estimated to have originated from a virus imported in 2015 and was most closely related to genomes from India. It is important to note that the implications of spread from India to Kenya are limited by sampling gaps, and India might not have been the direct source of the Kenyan outbreak. Indeed, the connection between the Somalian and Kenyan outbreaks indicates that the virus was most probably introduced from Somalia, and thus, that this strain is most likely spreading in African countries other than Kenya. Importantly, however, our results indicate that the *Ae. aegypti* adapted strain is no longer confined to India, but is spreading and causing large outbreaks in other countries and other continents of the world. Interestingly, all recent CHIKV ECSA lineage genomes from the *Ae. aegypti* prevalent regions of the Indian Ocean sub-lineage contain the two *Ae. aegypti* adaptive mutations, suggesting a possible replacement of the wild type by this enhanced fitness strain.

After the 2016 Kenyan outbreak resolved, a new CHIKV outbreak started in the coastal city of Mombasa in 2017-2018, with a large proportion of cases (70%) reporting severe joint pain and high fever (41). No publicly available genomes exist from this event as of yet, but given the proximity of the *Ae. aegypti* adaptive outbreak of 2016, and given this strain’s potential to cause large outbreaks, it is very possible that it is also responsible for this recent occurrence in Mombasa. Furthermore, other simultaneous CHIKV outbreaks have been reported in proximate regions to both Kenya and India, the 2016 outbreak in Mozambique and the 2017 outbreak in Dhaka, Bangladesh (42, 43). The virus from the Dhaka outbreak contained the K211E adaptive mutation, and despite the lack of E2 sequencing, our analyses of the E1 region placed it within the *Ae. aegypti* adapted cluster, suggesting that this outbreak was also caused by the adapted strain (43). Sequencing and analyzing complete viral genomes from these and other countries will aid in the tracking of the *Ae. aegypti* adaptive strain’s spread, and it will provide insight into its possible replacement of the wild type in the *Ae. aegypti* prevalent regions of the world. In addition, the association between *Ae. aegypti* competence and the E1:K211E and E2:A264V mutations should be investigated further, with competence studies utilizing regional mosquito populations. The level of increased receptiveness of these vectors to CHIKV infection may provide information for implementation of additional or alternate vector control strategies.

In conclusion, we show that the *Ae. aegypti* adaptive strain, carrying the E1:K211E and E2:A264V mutations, is capable of rapid spread in the *Ae. aegypti-*rich regions. It has recently caused several large CHIKV outbreaks in Africa and Asia, and it may have the potential to completely replace the wild type strain in these regions of the world resulting in widespread outbreaks around the globe.

## Disclaimer

Material has been reviewed by the Walter Reed Army Institute of Research. There is no objection to its presentation and/or publication. The opinions or assertions contained herein are the private views of the author, and are not to be construed as official, or as reflecting true views of the Department of the Army or the Department of Defense. The investigators have adhered to the policies for protection of human subjects as prescribed in AR 70–25.

## Supporting Information Legends

**S1_Checklist: STROBE Checklist**

**S2_Appendix. Infection confirmation, virus isolation and Sanger sequencing.** Detailed materials and methods used for sample collection, cell culture propagation, nucleic acid extraction, cDNA synthesis, PCR amplification, and Sanger sequencing confirmation.

